# A systems-level proteomic analysis identifies kinesin targets of KIFBP during neuronal development

**DOI:** 10.64898/2026.07.17.739260

**Authors:** Sarah-Catherine Paschall, Teresa Lynne Blasius, Alex Missman, Princess Rodriguez, Michael A. Cianfrocco, Kristen J. Verhey, Jason Stumpff

**Affiliations:** Department of Molecular Physiology and Biophysics, University of Vermont, Burlington, Vermont; Department of Cell and Developmental Biology, University of Michigan, Ann Arbor, Michigan; Life Sciences Institute, University of Michigan, Ann Arbor, Michigan; Department of Microbiology & Molecular Genetics, University of Vermont, Burlington, Vermont; Department of Biological Chemistry, University of Michigan, Ann Arbor, Michigan

## Abstract

Kinesins are molecular motor proteins essential for organizing and remodeling the cytoskeleton during neuronal development and maintenance. One key regulator is kinesin family binding protein (KIFBP), which inhibits a subset of kinesins by blocking motor-microtubule interactions. Homozygous mutations in KIFBP cause Goldberg-Shprintzen Syndrome (GOSHS), a neurodevelopmental disorder characterized by intellectual disability, microcephaly, and axonal neuropathy. Although loss of KIFBP has been linked to reduced neurite length and microtubule disorganization, the specific kinesins underlying these phenotypes remain unclear. Here we use a CRISPR-Cas9 generated KIFBP knockout Neuro-2a cell line to demonstrate that KIFBP is required for neurite extension and use inducible GFP-KIFBP to define the KIFBP interactome during neuronal differentiation. Immunoprecipitation coupled with mass spectrometry identified both known and novel KIFBP-associated kinesins. Single molecule TIRF microscopy confirmed direct inhibition of a subset of kinesins that co-immunoprecipitated with KIFBP. Notably, we identified KIF5A and KIF18B as previously unrecognized regulatory targets with potential roles in neuronal development. Together, these findings establish Neuro-2a cells as a model for studying KIFBP function and provide new insight into the regulation of kinesin activity and cytoskeletal dynamics in neurons.

## INTRODUCTION

The neuronal cytoskeleton is a dynamic and complex network that undergoes extensive remodeling during development. The polarized nature of neurons relies on precise organization of microtubules, which support axon and dendrite formation, enable intracellular transport, and facilitate synapse formation. Accordingly, microtubules and transport along them must be tightly regulated to ensure proper neuronal development and maintenance (Kapitein & Hoogenraad, 2015). Kinesin motors proteins are key contributors to this regulation, functioning in both cargo transport and modulation of microtubule dynamics (Hirokawa et al., 2010). For example, the kinesin-3 motor, KIF1A, transports synaptic vesicle precursors along the microtubules and down the axon, while the kinesin-8 motor, KIF18A, regulates microtubule dynamics (Hirokawa et al., 2010; Kevenaar et al., 2016; Okada et al., 1995). Despite these established roles, how kinesin activity is precisely controlled during neurodevelopment remains incompletely understood.

Multiple mechanisms regulate kinesin activity, including auto-inhibition, post-translational modification, and interaction with binding partners (Atherton et al., 2020; Verhey & Hammond, 2009). One such regulator, kinesin family binding protein (KIFBP), acts as a negative regulator of a subset of kinesins by binding and remodeling their motor domains to prevent motor-microtubule interactions (Atherton et al., 2020; Kevenaar et al., 2016; Solon et al., 2021).

Human genetic studies have linked KIFBP mutations to Goldberg-Shprintzen syndrome (GOSHS), a neurological disorder characterized by intellectual disability, microcephaly, and axonal neuropathy (Brooks et al., 2005; MacKenzie et al., 2020).

Consistent with this, loss of KIFBP function in animal models leads to reduced axon length and disorganized microtubules, underscoring its importance in neuronal development (Chang et al., 2019; Lyons et al., 2008). However, cellular studies have reported conflicting effects of KIFBP perturbation on neurite outgrowth (Drevillon et al., 2013; Kevenaar et al., 2016), highlighting an incomplete understanding of its function. Notably, the specific kinesins regulated by KIFBP in neurons, particularly during neurodevelopment, remain incompletely defined.

Here, we establish a Neuro-2a cell-based model to investigate KIFBP function during neuronal differentiation and to define the subset of kinesins it regulates. Using a CRISPR-Cas9 generated KIFBP knockout Neuro-2a cell line, we show that KIFBP is required for neurite extension. Affinity purification coupled with mass spectrometry identified 15 kinesins that associate with KIFBP, and single molecule analyses confirmed direct inhibition of a subset of these motors, including two previously unrecognized targets. Together, these findings establish a tractable cellular model system for studying KIFBP function and expand the subset of kinesins regulated by KIFBP during neuronal differentiation.

## RESULTS AND DISCUSSION

### Neuro-2a cells differentiate to form neurite extensions

The Neuro-2a mouse neuroblastoma cell line (N2A-A5) was used as a model to study neuronal differentiation (Dos Santos et al., 2023). Following 96 hours of serum reduction and retinoic acid treatment, N2A-A5 cells exhibited elongated neurite extensions compared to undifferentiated controls, consistent with previous studies (Kumar & Katyal, 2018) (Figure 1A). Quantitative immunofluorescent microscopy revealed a significant increase in both neurite length and the percentage of cells with neurite outgrowths (differentiated cell percent) after 96 hours (Figure 1B-C).

**Figure 1.**
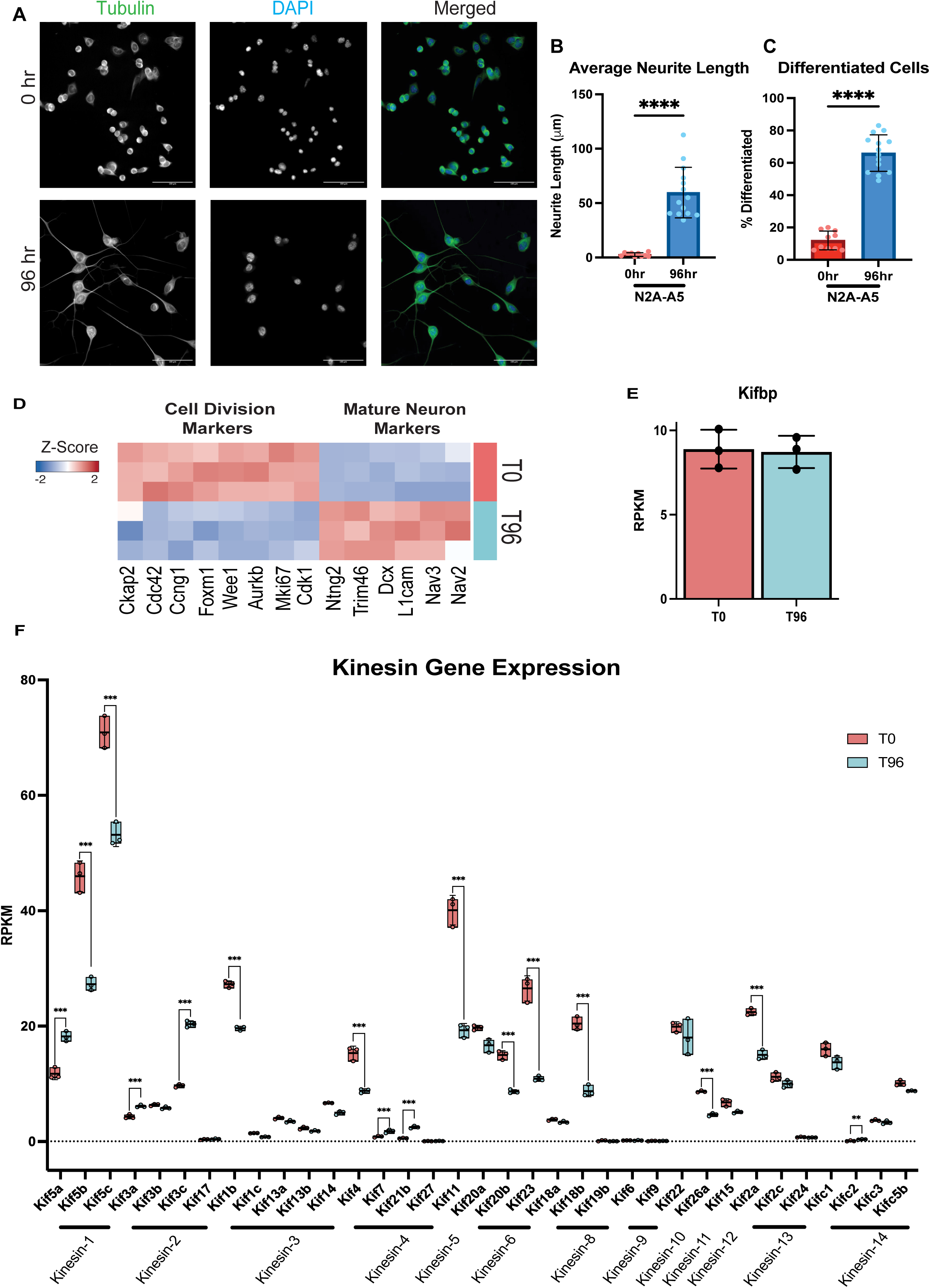
N2A-A5 cells differentiate into neuron-like cells and express 35 kinesin proteins. **(A)** Undifferentiated N2A-A5 cells were fixed at 0 hours before differentiation treatment. Differentiated N2A-A5 cells fixed after 96 hour differentiation treatment (1% FBS and 10 µM retinoic acid). Colors indicate pseudo-color in merged images. Brightness/contrast levels are set differently to optimize visualization. Scale bar is 100 µm. **(B)** Quantification of average neurite length in differentiated N2A-A5 cells. **(C)** Quantification of the percent of differentiated cells in each population. For (**B, C**) data were analyzed via an unpaired t-test with Welch’s correction. P value style: <0.0001 (****). RNA sequencing was performed on N2A-A5 cells at time 0 hour (T0) and differentiated cells after 96 hours (T96). **(D)** Gene markers for mature neurons and cell proliferation are altered between time points (FDR < 0.05). **(E)** Barplot of Kifbp expression between T0 and T96. **(F)** Box plots displaying expression of 35 kinesins between T0 and T96 organized by kinesin family. For (**E, F**) expression in reads per kilobase per million (RPKM) normalized expression for cell types. Error bars represent S.D. and dots represent individual data points. Data represents three independent samples for 0 hour and 96 hour groups. FDR: < 0.001 (***), < 0.01 (**), <0.05 (*). If no significance is shown result was not significant (> 0.05).

### Differentiated N2A-A5 cells express mature neuron markers and kinesin motor proteins

To further validate N2A-A5 cells as a model system, bulk RNA sequencing was performed to investigate transcriptional programs of undifferentiated (0 hr) and differentiated (96 hr) cells. Comparison between N2A-A5 (96 hr vs. 0 hr) cells revealed 1,296 upregulated and 448 downregulated differentially expressed genes (DEG absolute log_2_FC>1 and FDR < 0.05; Figure S1A-B). Principal component analysis (PCA) revealed that differentiation was the major source of variation (Figure S1C). Pathway analysis of DEGs revealed many DEGs mapped to extracellular matrix pathways (Figure S1D). Examples of genes altered include mature neuron markers such as L1cam and Dcx, involved in neurite outgrowth and neurogenesis, and common cell proliferation markers such as Cdk1 and Mki67 (FDR < 0.05). (Alberts et al., 2003; Couillard-Despres et al., 2005; Massacci et al., 2023; Whitfield et al., 2006) (Figure 1D). Furthermore, we found that Kifbp expression remains steady following differentiation. (Figure 1E).

Since KIFBP is a known binding partner for a subset of kinesins, we investigated kinesin gene expression levels in these samples. Our results indicate that 35 different kinesin genes are expressed in N2A-A5 cells (Figure 1F). Importantly, kinesins with critical roles in neuronal development and maintenance, such as Kif5a and Kif3a, are significantly up-regulated in differentiated N2A-A5 cells (Hirokawa et al., 2010; Nishimura et al., 2004; Xia et al., 2003) (Figure 1F). However, two kinesins involved in neuronal transport, Kif1c and Kif1a, displayed reduced or undetectable expression in differentiated cells, respectively. (Figure 1F) (Hirokawa et al., 2010; Okada et al., 1995; Schlager et al., 2010). Taken together, these results suggest that differentiated N2A-A5 cells are a useful model to study kinesin regulation by KIFBP.

### Loss of KIFBP impairs neurite extension

To assess how loss of KIFBP affects neurite formation in N2A-A5 cells, two independent KIFBP knockout cell lines were generated using CRISPR-Cas9 genome editing. As a control, N2A-A5 cells were treated with the plasmid containing the Cas9 enzyme without targeted guide RNA. Sequencing of genomic DNA from both knockout clones confirmed frameshift mutations in exon 2 resulting in early stop codons, while the control clone contained wild type KIFBP sequence. Immunoblotting of both undifferentiated and differentiated knockout cell lines verified loss of KIFBP protein (Figure 2A, Figure S2A). Following differentiation, quantitative immunofluorescent microscopy revealed that KIFBP knockout cells exhibited significantly shorter neurites compared to control cells (Figure 2B-C). In addition, loss of KIFBP also decreases the percent of differentiated cells with neurite outgrowths compared to control cells (Figure 2D). These findings indicate that KIFBP is required for efficient neurite extension in N2A-A5 cells, consistent with its requirement for establishing proper neuron length in animal models (Chang et al., 2019; Lyons et al., 2008).

**Figure 2.**
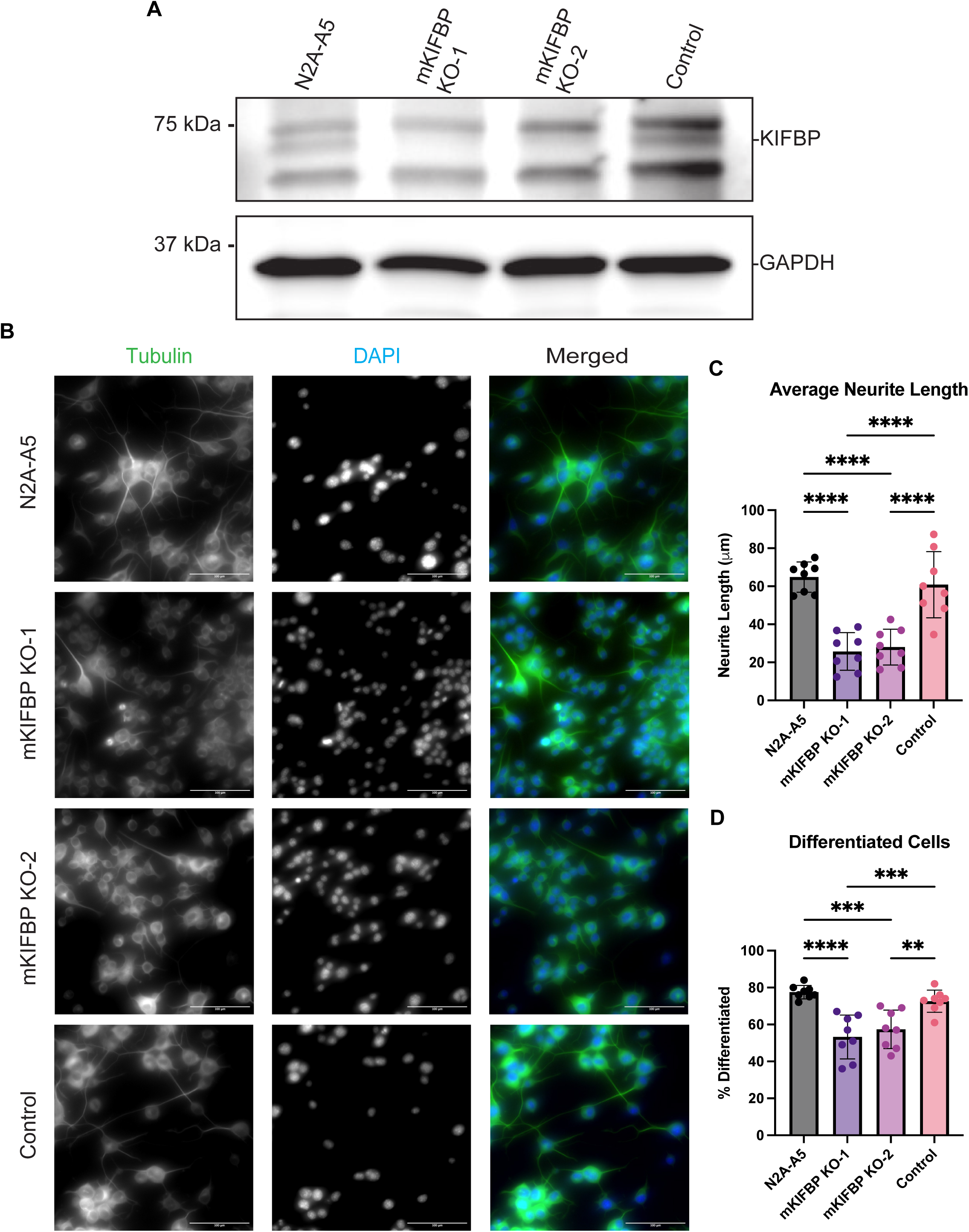
KIFBP knockout cell lines display neurite length defects. **(A)** Western blot of KIFBP, showing protein levels expressed in N2A-A5, KIFBP knockout clone #1, KIFBP knockout clone #2, and N2A-A5 control cells. GAPDH was used as a loading control. **(B)** Representative immunofluorescence images of N2A-A5, N2A-A5 KIFBP knockout cell lines 1 and 2, and N2A-A5 control cells after differentiation treatment for 96 hours. Colors indicate pseudo-color in merged images. Brightness/contrast levels are set differently across cell lines to optimize visualization. Scale bar is 100 µm. **(C)** Quantification of average neurite length in differentiated N2A-A5, N2A-A5 KIFBP knockout cell lines 1 and 2, and N2A-A5 control cell lines. **(D)** Quantification of the percent of differentiated cells in each population. For (**C, D**) data were analyzed via a one-way ANOVA with Tukey’s test for multiple comparisons. P value style: <0.0001 (****), <0.001(***), <0.01(**). If no significance is indicated, the result was not significant (>0.05).

### KIFBP associates with a subset of kinesins during neuronal differentiation

Our results suggest that KIFBP is important for neuronal cell differentiation and proper neurite formation. KIFBP-kinesin interactions were previously identified using HEK293 and HeLa cells, however, the KIFBP interactome during neurodevelopment has not been defined (Kevenaar et al., 2016). To address this gap, N2A-A5 cells were generated to express GFP-tagged wild type mouse KIFBP under a doxycycline inducible promoter. Live cell imaging was performed after differentiation to validate the expression of GFP or GFP- mKIFBP constructs (Figure 3A). Notably, our results show that loss of KIFBP impaired neurite growth (Figure 2C), whereas overexpression of GFP or GFP-mKIFBP did not alter neurite length or differentiation efficiency, indicating this system does not perturb neuronal morphology (Figure 3B-D). However, previous studies have reported conflicting results regarding the effects of KIFBP overexpression on neurite length, with increased length being reported in SH-SY5Y cells and reduced length reported in hippocampal neurons (Drevillon et al., 2013; Kevenaar et al., 2016). This suggests that the optimal level of KIFBP expression is needed to control the precise length of neurons during neuronal development and that this may vary by cell type.

**Figure 3.**
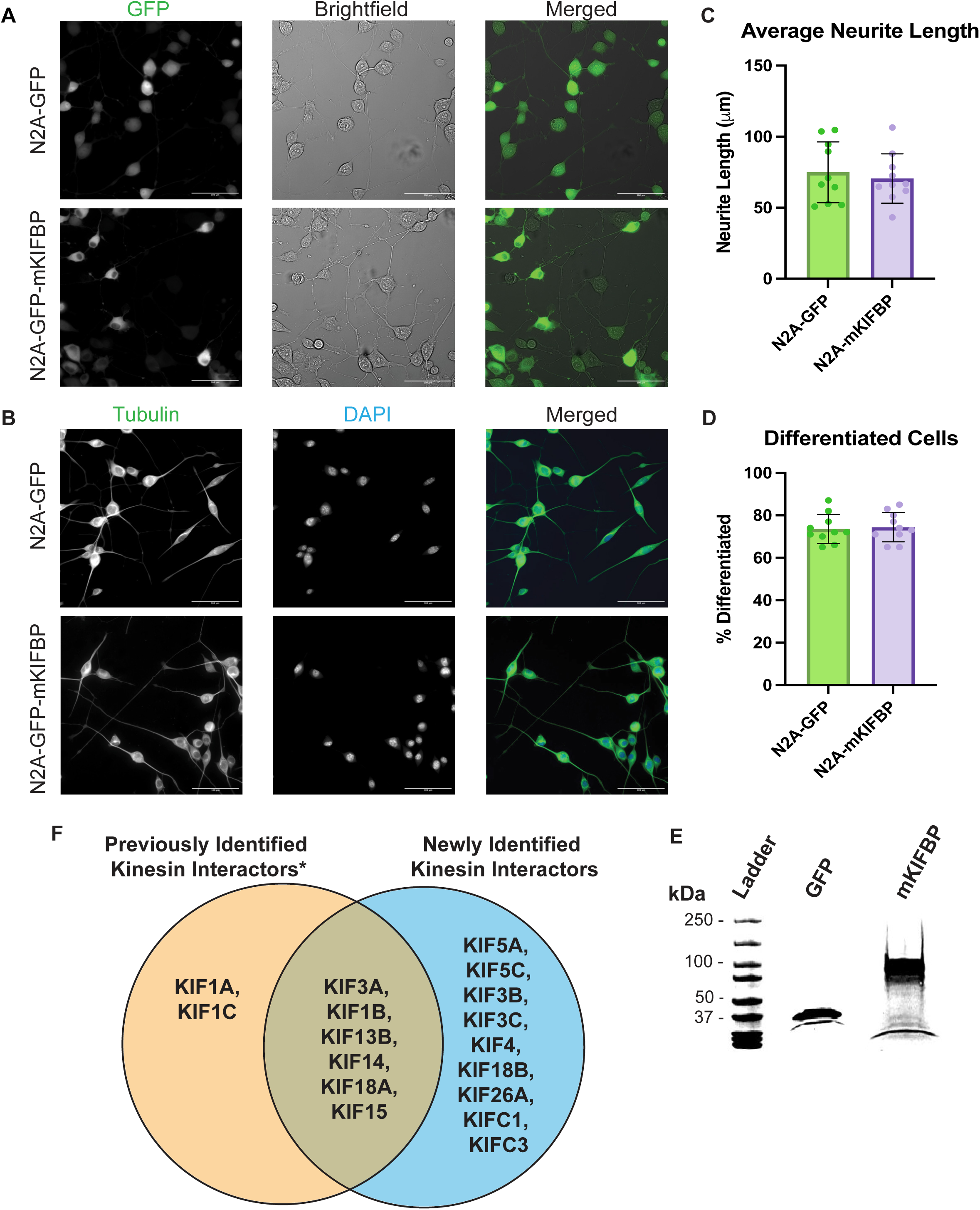
KIFBP binds a subset of kinesins in differentiated N2A-A5 cells. **(A)** Representative live cell images of differentiated N2A-A5 cells expressing GFP or GFP-mKIFBP. Colors indicate pseudo-color in merged images. Brightness/contrast levels are set differently to optimize visualization. Scale bar is 100 µm. **(B)** Representative immunofluorescent images of N2A-A5 cells expressing GFP or GFP-mKIFBP after differentiation treatment for 96 hours. Colors indicate pseudo-color in merged images. Brightness/contrast levels are set differently to optimize visualization. Scale bar is 100 µm. **(C)** Quantification of average neurite length in differentiated N2A-A5 GFP or GFP-mKIFBP cell lines. **(D)** Quantification of the percent of differentiated cells in each population. For (**C, D**) data were analyzed via an unpaired t-test with Welch’s correction. If no significance is indicated result was not significant (> 0.05). **(E)** Coomassie-blue stained SDS-PAGE gel of immunoprecipitation samples from differentiated N2A-A5 cells expressing GFP alone or GFP-mKIFBP. GFP sample expressed at 37 kDa and GFP-mKIFBP sample expressed at 100 kDa. **(F)** Previously reported kinesin binding partners of KIFBP (* (Kevenaar et al., 2016)) compared to newly reported kinesin binding partners of KIFBP. Kinesin proteins were considered a KIFBP interactor if the kinesin was reported in at least 2 mass spectrometry protein analyses with more than 1 reported unique peptide.

Affinity purification followed by mass spectrometry identified 338 KIFBP-interacting proteins, including 15 kinesin proteins (Figure 3E-F and supplemental data 1). Several of these kinesin motors were previously reported to interact with KIFBP including KIF1B, KIF3A, and KIF18A (Kevenaar et al., 2016; Solon et al., 2021; Wozniak et al., 2005) (Figure 3F). We also identified new kinesin interactors including KIF3B, KIF18B, and KIF26A (Figure 3F). A prior study identified binding interactions of KIFBP with the kinesin-3s, KIF1A and KIF1C and demonstrated that KIFBP blocks their motility in cellular transport assays (Kevenaar et al., 2016) (Table I). The absence of these interactions in our IP-MS analysis is likely explained by their low or absent expression in differentiated N2A-A5 cells determined by our RNA sequencing results, highlighting the context dependence of KIFBP-kinesin interactions. Notably, previously reported interactions with stathmin-like protein (STMN2) and the charged multivesicular body protein 2b (CHMP2b) were not observed despite detectable mRNA expression in N2A-A5 cells (Alves et al., 2010; Kirwan et al., 2025) (Figure S1E-F).

### KIFBP directly inhibits a subset of kinesin partners identified in N2A-A5 cells

To determine which of the kinesins identified in our IP-MS analysis (Figure 3) are directly regulated by KIFBP, we performed single-molecule motility assays using total internal reflection fluorescence (TIRF) microscopy. We generated a constitutively-active version of each kinesin of interest by fusing the minimal motor domain to mNeonGreen (mNG). The tagged motor domains were expressed in COS-7 cells and cell lysates were incubated with purified KIFBP protein or buffer and then added to flow cells containing taxol-stabilized microtubules.

We first tested whether KIFBP inhibits the motility of the kinesin-1 members KIF5A and KIF5C. KIF5C-mNG was observed to undergo short periods of directed movement along microtubules, and this behavior was not altered in the presence of KIFBP (Figure 4A). In contrast, KIF5A-mNG was observed to undergo long directed runs along microtubules, and this motility was dramatically reduced in the presence of KIFBP (Figure 4A). This result is particularly notable, as other members of the kinesin-1 family, such as KIF5B and KIF5C, do not interact with KIFBP (Kevenaar et al., 2016) (Figure 4A). This unexpected specificity suggests that KIFBP can discriminate among closely related motors and may preferentially regulate neuronally enriched kinesins. Given the established role that KIF5A plays in anterograde axonal transport of organelles and cargo its inhibition by KIFBP suggests a mechanism for coordinating cargo transport during neuronal (Cozzi et al., 2024; Kumar et al., 2023).

**Figure 4.**
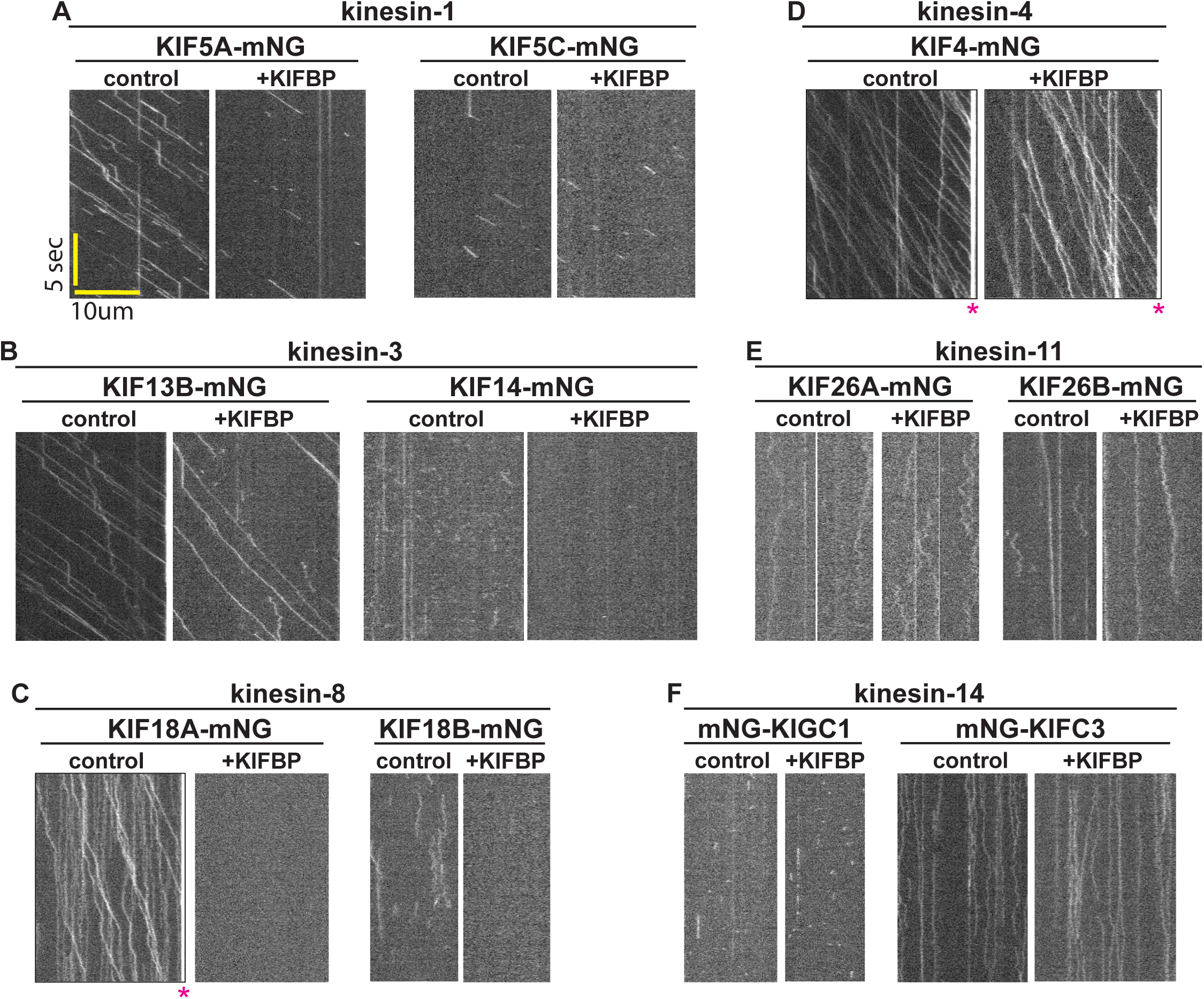
KIFBP directly inhibits a subset of kinesins involved in neuronal differentiation. **(A-F)** Representative kymographs of single molecule motility assays. The kinesin of interest was expressed in COS-7 cells and added to a flow chamber in a cell lysate and in the presence of buffer control or purified KIFBP protein. Time is on the x-axis, distance is on the y-axis. Microtubule tip accumulation represented by (*). Single-molecule behavior in the absence or presence of KIFBP was determined for **(A)** the kinesin-1 members KIF5C and KIF5C, **(B)** the kinesin-3 members KIF13B and KIF14, **(C)** the kinesin-8 members KIF18A and KIF18B, **(D)** the kinesin-4 member KIF4, **(E)** the kinesin-11 members KIF26A and KIF26B, and **(F)** the kinesin-14 members KIFC1 and KIFC3.

Interestingly, our mass spectrometry data reported that KIFBP interacted with KIF3A, KIF3B, and KIF3C. These kinesin-2 family motors generally exist in heterodimeric motor complexes (Hirokawa et al., 2009; Muresan et al., 1998). We were unable to test the effects of KIFBP on motility of the kinesin-2 family members KIF3A/KIF3B and KIF3A/KIF3C due to difficulties in generating motile, dimeric versions of these proteins. However, our binding assays, structural analysis, and cellular assays indicate that KIFBP binds directly to and inhibits the motility of KIF3A, KIF3B, and KIF3C (Missman et al, manuscript in preparation). This was surprising as previous co-IP experiments suggested that KIFBP binds to KIF3A and KIF3C but not to KIF3B (Kevenaar et al., 2016).

The kinesin-3 family member KIF13B-mNG undergoes long directed runs along microtubules, and this motility was dramatically reduced in the presence of KIFBP (Figure 4B). KIF14-mNG undergoes transient diffusive interactions with microtubules, and these events were also dramatically reduced by KIFBP (Figure 4B). The kinesin-8 family member KIF18A-mNG undergoes long directed runs and accumulates at the plus ends of the microtubules, whereas KIF18B-mNG undergoes diffusive movement along the microtubules. Both motors were inhibited by KIFBP (Figure 4C). That KIFBP binds to and inhibits KIF13B, KIF14, and KIF18A is consistent with previous work (Atherton et al., 2020; Kevenaar et al., 2016; Solon et al., 2021). However, the ability of KIFBP to bind to and inhibit KIF18B was unexpected, as no interaction was detected between these proteins by co-IP (Kevenaar et al., 2016). KIFBP thus appears to be a general inhibitor of both the kinesin-3 and kinesin-8 families.

Finally, this work identified several kinesin proteins by IP-MS analysis which KIFBP was unable to inhibit in our single-molecule assays. Specifically, the kinesin-4 KIF4-mNG undergoes long directed runs and accumulates at the plus ends of microtubules, but neither behavior was inhibited by KIFBP (Figure 4D). The kinesin-11 proteins, KIF26A-mNG and KIF26B-mNG, undergo diffusive movement along microtubules but this behavior was not inhibited by KIFBP (Figure 4E). The kinesin-14s, KIFC1-mNG and KIFC3-mNG, also displayed diffuse behavior on microtubules and were not inhibited by KIFBP (Figure 4F).

Together, these results support the previously suggested model that KIFBP regulates neuronal development by selectively inhibiting kinesins and expands the subset of kinesins known to be directly regulated by KIFBP (Table 1). This expanded interaction network provides a framework for understanding how dysregulation of kinesin activity may contribute to Goldberg-Shprintzen syndrome. Future studies should determine how individual KIFBP-kinesin interactions contribute to neurodevelopment and disease (Brooks et al., 2005; MacKenzie et al., 2020).

**Table 1.** Summary table of KIFBP-kinesin binding partners.

| Family | Kinesin | This Study |  | Previous Work |  |  | Known Role In Neurons |
| --- | --- | --- | --- | --- | --- | --- | --- |
|  |  | IP/MS/In Vitro Pull Down | Single Molecule Inhibition | IP/MS/In Vitro Pull Down | Co-IP | Cellular Transport Inhibition |  |
| Kinesin-1 | KIF5A | + | + |  |  |  | Transport (Xia et al., 2003) |
|  | KIF5B | - | N.D. | -<sup>2</sup> | -<sup>2</sup> | -<sup>2</sup> | Transport (Tanaka et al., 1998) |
|  | KIF5C | + | - | -<sup>2</sup> | -<sup>2</sup> |  | Maintenance (Kanai et al., 2000) |
| Kinesin-2 | KIF3A | + | (+) | +<sup>2</sup> | +<sup>2</sup> | +<sup>2</sup> | Transport (Nishimura et al., 2004) |
|  | KIF3B | + | (+) | -<sup>2</sup> | -<sup>2</sup> |  | Transport (Yamazaki et al., 1995) |
|  | KIF3C | + | (+) | -<sup>2</sup> | -<sup>2</sup> |  | MT Regulation (Gumy et al., 2013) |
| Kinesin-3 | KIF1A | - | N.D. | +<sup>1,2</sup> | +<sup>1</sup> | +<sup>1,2</sup> | Transport (Okada et al., 1995) |
|  | KIF1B | + | N.D. | +<sup>1,2</sup> | +<sup>1</sup> | +<sup>1,2</sup> | Transport (Nangaku et al., 1994) |
|  | KIF1C | - | N.D. | +<sup>1,2</sup> | +<sup>1</sup> | +<sup>1,2</sup> | Transport (Schlager et al., 2010) |
|  | KIF13B | + | + | +<sup>1,2</sup> | +<sup>1</sup> | +<sup>2</sup> | Transport (Horiguchi et al., 2006) |
|  | KIF14 | + | + | +<sup>1,2</sup> | +<sup>1</sup> |  | Maintenance (Fujikura et al., 2013) |
| Kinesin-4 | KIF4 | + | - | -<sup>2</sup> | -<sup>2</sup> |  | Maintenance (Sekine et al., 1994) |
| Kinesin-8 | KIF18A | + | + | +<sup>1,2,3</sup> | +<sup>1,3</sup> | +<sup>1,2</sup> | MT Regulation (Kevenaar et al., 2016) |
|  | KIF18B | + | + | -<sup>1</sup> | -<sup>1</sup> |  | Unknown |
| Kinesin-11 | KIF26A | + | - |  |  |  | MT Regulation (Qian et al., 2022) |
| Kinesin-12 | KIF15 | + | N.D. | +<sup>1,2,3</sup> | +<sup>1,3</sup> | +<sup>1,2</sup> | MT Regulation (Dong et al., 2019) |
| Kinesin-14 | KIFC1 | + | - |  |  |  | MT Regulation (Muralidharan et al., 2022) |
|  | KIFC3 | + | - |  |  |  | MT Regulation (Cao et al., 2020) |

Key: (+); Missman et al., manuscript in preparation. N.D.; Not Done. <sup>1</sup>; (Atherton et al., 2020). <sup>2</sup>; (Kevenaar et al., 2016). <sup>3</sup>; (Solon et al., 2021)
KIFBP Directly Inhibits Kinesin-MT Binding
KIFBP Does Not Inhibit Kinesin-MT Binding

## MATERIALS AND METHODS

### Cell Culture and Differentiation

N2A-A5 acceptor cells were gifted by Eugene Makeyev, Kings College, London (Khandelia et al., 2011). N2A-A5 acceptor cells were cultured at 37°C with 5% CO_2_ in Dulbecco’s Modified Eagle Medium (DMEM; Gibco, 11965118) supplemented with 10% fetal bovine serum (FBS; Gibco 16000044), 1% Penicillin Streptomycin (Pen/Strep; Gibco 15140122), 1% Sodium Pyruvate (Gibco, 11360070), and 10 µg/mL blasticidin (Thermo Fisher, R21001). N2A-A5 cells were differentiated for 96 hours by seeding in differentiation culture media without blasticidin, supplemented with 1% FBS and 10 µM retinoic acid (MedChem, HY14649).

### RNA Sequencing Sample Preparation

Total RNA was extracted from undifferentiated (0 hour) and differentiated (96 hour) N2A-A5 cells using the RNeasy Plus Micro Kit (Qiagen, 74034). After extraction, RNA was stored in RNase-free water supplemented with 40 U/uL RiboLock RNase Inhibitor (Thermo Scientific, EO0381) at -80°C.

### Bulk RNA-Seq library preparation and sequencing

To quantify gene-level expression changes in undifferentiated (0 hr) and differentiated (96 hr) N2A-A5 cells, we performed bulk RNA sequencing (RNA-seq; n=3). Total RNA was extracted using the RNeasy Plus Micro Kit (Qiagen) and RNA integrity was assessed (RIN ≥ 7) with the Agilent Bioanalyzer. Ribosomal RNA was depleted and libraries were prepared using the Takara SMARTer Stranded Total RNA-Seq Kit v2 – Pico Mammalian. Libraries were sequenced on the Singular Genomics G4 platform to generate pair-end 150 bp reads to a target depth of ∼30 million reads per sample.

### Bulk RNA-Seq analysis

Raw sequencing data was processed using the nf-core/rnaseq pipeline (Ewels et al., 2020). Raw reads were assessed with FASTQC and aligned to the mm39 reference genome using both STAR and Salmon to enable complementary gene- and transcript-level quantification (Andrew, 2010; Dobin et al., 2013; Patro et al., 2017). Differential gene expression analysis was performed using DESeq2 (Love et al., 2014). Multiple hypothesis testing was controlled using the Benjamini–Hochberg false discovery rate (FDR), and genes were considered significantly differentially expressed (DEGs) if they meet both an adjusted p-value threshold (FDR < 0.05) and an absolute log2 fold change ≥ 1.

### Generation and Validation of KIFBP Knockout Cell Lines

The full-length sequence of the mouse KIFBP gene was used to design sgRNAs targeting exon 1 and exon 2 (NCBI Gene ID: 72320). Each sgRNA was generated into the pSPCas9 BB-2A-Puro (PX549) v2.0 plasmid by GenScript BioTech. The resulting plasmid was transfected into N2A-A5 cells with recombinase using Lipofectamine LTX transfection (Invitrogen; 15338030). The pSPCas9 BB-2A-Puro (PX549) v2.0 plasmid without sgRNA was used to generate N2A-A5 control cells. 24 hours after LTX transfections, N2A-A5 cells that had undergone homologous recombination were selected with 2.5 µg/mL puromycin for 24 hours, followed by selection with 5 µg/mL puromycin for 48 hours. Single cells were then isolated in 96 well plates (Falcon; 353072) by serial dilution and allowed to expand until wells were 50% confluent. After single clone expansion, genomic DNA was extracted using the QIAmp DNA Blood Mini Kit (Qiagen; 51104) and amplified by PCR. DNA bands at the predicted size were isolated through gel extraction (Qiagen; 28706) and subject to sequencing (Eurofins).

### Western Blotting

For KIFBP western blot detection, 1x10^6^ cells were collected and lysed in RIPA Lysis and Extraction Buffer (Thermo Scientific; 89901) with 1X Halt Protease and Phosphatase Inhibitor (Thermo Scientific; 78440) by rotating at 4°C for 30 minutes. Cell lysates were spun at 14,000 xg for 10 minutes and the resulting supernatant was collected. Total protein concentration was determined using Pierce Dilution-Free Rapid Gold BCA Protein Assay (Thermo Scientific; A55862). The remaining supernatant was added to 4X Laemmli sample buffer (BioRad; 1610747) prepared with 2-mercaptoethanol (BioRad; 1610710). Equal concentrations of samples were then loaded into 4-15% tris-glycine polyacrylamide gels (Bio-Rad; 4561083), separated by electrophoresis, and transferred to Immuno-Blot low fluorescence polyvinylidene fluoride membranes (Bio-Rad; 1620261). Membranes were blocked in 5% TBST blotting grade blocker (Bio-Rad; 1706404) and blotted using rabbit anti- KIFBP (1:500, ProteinTech; 25653-1-AP) and rabbit anti-GAPDH (1:1000, Invitrogen; PA1- 987) primary antibodies incubated overnight at 4°C. Blots were incubated with donkey anti-rabbit HRP (1:1000, Thermo Scientific; A16035) secondary antibodies at room temperature for 1 hour. Imaging was preformed using Clarity Max Western ECL Substrate (Bio-Rad; 1705062) on a Bio-Rad ChemiDoc and images were processed using ImageJ/FIJI.

### Generation and Validation of N2A-A5 Inducible Cell Lines

Inducible cell lines were generated using recombination-mediated cassette exchange as previously described (Khandelia et al., 2011). Briefly, a wild type mouse KIFBP puromycin resistant plasmid containing LoxP sites was generated by GenScript BioTech. The resulting plasmid was transfected into N2A-A5 cells with recombinase using Lipofectamine LTX transfection (Invitrogen; 15338030). 24 hours after LTX transfections, N2A-A5 cells that had undergone exchange were selected with 2.5 µg/mL puromycin for 48 hours, followed by selection with 5 µg/mL puromycin for 72 hours. Cells were then maintained in culture media with 2.5 µg/mL puromycin for continued selection. To confirm the sequence of inserted DNA in the selected cell populations, genomic DNA was extracted using the QIAmp DNA Blood Mini Kit (Qiagen; 51104) and amplified by PCR. DNA bands at the predicted size were isolated through gel extraction (Qiagen; 28706) and subject to sequencing (Eurofins). Expression of inserted DNA sequences was induced via treatment in normal N2A-A5 growth media or differentiation media with 2 µg/mL doxycycline (Fisher Scientific; BP26531).

### Immunoprecipitation

N2A-A5 inducible cell lines were seeded in 150 mm cell culture dishes (Falcon; 353025) and subject to differentiation treatment for 96 hours in differentiation media with 2 µg/mL doxycycline to induce expression of GFP constructs. Differentiated N2A-A5 inducible cell lines were collected and lysed in ice cold 10 mM tris buffer, pH 7.5, with 150 mM NaCl, 0.5 mM EDTA, 0.5% Triton X-100 (Thermo Scientific; A16046AE), and 1X Halt Protease and Phosphatase Inhibitor (Thermo Scientific; 78440) by rotating at 4°C for 30 minutes. GFP immunoprecipitation was performed using GFP-Trap nanobody coated magnetic particles (ChromoTek; GTD-20) according to the manufacturer’s instructions. Samples were separated by electrophoresis on 4-15% tris-glycine polyacrylamide gels (Bio-Rad; 4561084) and stained overnight with Coomassie Brilliant Blue (Bio-Rad; 1610400). After de-staining, samples were extracted and stored in LC/MS grade water (Fisher Scientific; W6500) at 4°C.

### LC-MS/MS and Data Analysis

Peptide separations were performed using an EASY-nLC 1200 UHPLC system (Thermo Fisher Scientific) equipped with an Acclaim PepMap 100 trap column (75 μm × 2 cm, C18, 3 μm, 100 Å; Thermo Fisher Scientific) and a DNV PepMap Neo analytical column (75 μm × 15 cm, C18, 2 μm, 100 Å; Thermo Fisher Scientific). The nanospray electrospray ionization (ESI) source was coupled to an Orbitrap Exploris 240 mass spectrometer (Thermo Fisher Scientific). Full MS spectra were acquired in the Orbitrap, and selected precursor ions were fragmented by higher-energy collisional dissociation (HCD) to generate MS/MS spectra.

Mobile phase A consisted of 0.1% formic acid (FA) in water, whereas mobile phase B consisted of 0.1% FA in 80% acetonitrile (Fisher Chemical). SpeedVac-dried tryptic peptides were resuspended in 15 μL of 0.1% FA containing 2.5% acetonitrile, and 5 μL of each sample was loaded onto the trap column. Peptides were separated on the analytical column using a linear gradient of 5–25% solvent B over 40 min, followed by 25–40% solvent B over 20 min, at a flow rate of 200 nL/min.

Mass spectrometry data were acquired in data-dependent acquisition (DDA) mode with a 2 s cycle time. Full MS scans were acquired over an m/z range of 375–1400 at a resolution of 60,000 with a normalized AGC target of 300%, automatic maximum injection time, and profile mode. MS/MS spectra were acquired at a resolution of 15,000 using an isolation window of 2 m/z, a standard AGC target, automatic maximum injection time, and a normalized collision energy (NCE) of 30%.

Raw data were searched against the UniProt mouse reference proteome database (UP000000589) using Proteome Discoverer 3.1 (Thermo Fisher Scientific). Search parameters included a precursor mass tolerance of 10 ppm and a fragment mass tolerance of 0.02 Da. Carbamidomethylation of cysteine residues (+57.021 Da) was specified as a fixed modification, whereas oxidation of methionine (+15.995 Da) and phosphorylation of serine, threonine, and tyrosine residues (+79.966 Da) were specified as variable modifications. Up to two missed tryptic cleavages and a maximum of three variable modifications per peptide were permitted. Peptide-spectrum matches were filtered using the Percolator algorithm to achieve a false discovery rate (FDR) of less than 1% at the peptide level.

### Immunofluorescence

For fixed cell immunofluorescence experiments, cells were seeded on acid-treated 12 mm glass coverslips in a 24 well dish and fixed in -20°C methanol with 1% paraformaldehyde (Electron Microscopy Sciences; 15710) for 10 minutes on ice. After fixation, coverslips were washed 3 times for 5 minutes each in Tris-Buffered Saline (1X TBS; 150 mM NaCl, 50 mM Tris base, pH 7.4) and then incubated with mouse anti-tubulin, clone DM1A, Alexa Fluor 488 conjugate (1:100, Sigma-Aldrich; 16232) diluted in antibody dilution buffer (AbDil; 1% bovine serum albumin (Sigma-Aldrich; A9418), 0.1% Triton X-100 (Thermo Scientific; A16046AE), 0.02% sodium azide (Fisher Scientific; BP922I) in TBS) for 2 hours at room temperature on an orbital shaker. After antibody incubation, coverslips were washed 2 times for 5 minutes each in 1X TBS, then washed once in 1X TBS with 125 ng/mL DAPI (Invitrogen; D1306), followed by 2 additional washes each for 5 minutes in 1X TBS.

Coverslips were mounted either on slides (ThermoFisher; 3011002) or in a 24 well glass bottom plate (Cellvis; P24-1.5H-N) using Prolong Gold mounting medium (Invitrogen; P36934).

### Plasmids

*Hs*KIF5A(1-560)-mNG, *Rn*KIF5C(1-560)-mNG, *Hs*KIF13B(1-412,ΔP391)-mNG, *Hs*KIF14(356- 737)-mNG, *Hs*KIF4(1-367)-LZ-mNG-TwinStrep, *Hs*KIF18A(1-452)-mNG, *Hs*KIF18B(1-395)- mNG, *Hs*KIF26A(369-739)-mNG, *Hs*KIF26B(448-815)-mNG, mNG-*Hs*KIFC1(145-673), mNG-*Hs*KIFC3(375-834) were generated in the pEGFP-N1 vector by traditional cloning or Gibson cloning.

### COS-7 Cell Lysates

COS-7 cells were grown in DMEM (Gibco; 11960044) with 10% Fetal Clone III serum (HyClone; SH30109.03) and 1% Glutamax (Gibco; 35050061). The cells were seeded into 6- well plates at 1x10^5^ cells per well. The following day, 1µg of plasmid DNA was transfected per well with Mirus Trans-IT LT1 (Mirus Bio; MIR2305) and Opti-MEM (Gibco; 31985062) according to the manufacturer’s instructions. Cells were harvested 24 hours post- transfection by treating with 0.05% Trypsin-EDTA (Gibco; 25300054) and then the trypsin was diluted with an equal volume of serum-containing medium. The cells were pelleted at 1,500 xg for 5 minutes at 4°C, washed with 1 mL of serum containing medium, and the cell pellets were aspirated completely and resuspended in 25 µl of ice-cold lysis buffer (25 mM Hepes/KOH, 115 mM potassium acetate, 5 mM sodium acetate, 5 mM MgCl_2_, 0.5 mM EGTA, and 1% Triton X-100, pH 7.4) freshly supplemented with 10 mM ATP, 1 mM PMSF, and protease inhibitors (Sigma; P8340). The lysate was clarified by centrifugation at 20,000 x g for 10 minutes at 4°C. Supernatants were aliquoted, flash-frozen in liquid nitrogen, and stored at -80°C until further use.

### Single-Molecule Motility Assays by Total Internal Reflection Fluorescence Microscopy

Flow chambers were assembled as described previously (Henty-Ridilla, 2022). Briefly, passivated #1.5 coverslips were attached to a glass slide by melting two strips of parafilm to act as a glue along opposite sides of the coverslip. Approximate chamber volume is 20 µl. TIRF assays were performed at room temperature on an inverted Nikon Ti-E/B total internal reflection fluorescence (TIRF) microscope with a perfect focus system, a 100 x 1.49 NA oil immersion TIRF objective, two 20 mW diode lasers (488 and 640 nm) and EMCCD camera (iXon^+^ DU879; Andor). Image acquisition was controlled using Nikon Elements software (Nikon Instruments).

Biotin/647 microtubules were polymerized up to one week before imaging. Microtubules were polymerized with 80 mg of unlabeled porcine brain tubulin (Cytoskeleton Inc.; T240), 10 µg of biotin tubulin (Cytoskeleton Inc.; T333P), and 5 µg of fluorescent HiLyte 647 tubulin (Cytoskeleton Inc.; TL670M) in BRB80 buffer supplemented with 2.5 mM GTP and 4 mM MgCl_2_ at 37°C for 30 minutes. A 6.7X volume of prewarmed BRB80 buffer containing 20 µM Paclitaxel (Cytoskeleton Inc.; TXD01) was then added to the microtubule mixture and maintained at 37°C for an additional 3 hours. Microtubules were stored at room temperature in the dark. Prior to using, microtubules were pelleted at 21,130 xg for 10 minutes at room temperature. The supernatant was gently aspirated and 10 µM Paclitaxel/BRB80 was added, incubated at 37°C for 2 minutes, and then gently mixed to resuspend. The day of use, a working stock was created by diluting microtubules 1:30 in 10 µM Paclitaxel/BRB80 and held at 37°C.

Flow chambers were sequentillary incubated as follows: Flow in 1 mg/mL biotin-BSA (Sigma; A8549) then wash with 1 mg/mL BSA/BRB80. Flow in 0.5 mg/mL NeutrAvidin/BRB80 (Thermo; 31000) then wash with 1 mg/mL BSA/BRB80. Flow in biotin/647 microtubules and incubate the chamber at 37°C for 3 minutes. During the incubation, assemble the motility mix: 30 µl of 15 µM Paclitaxel/P12 (P12: 12 mM PIPES/KOH pH 6.8, 2 mM MgCl_2_, 1 mM EGTA), 10 µl of 30 mg/mL BSA/P12, 2 µl of 10 mg/mL casein (Sigma; C8654)/P12, 3 µl of oxygen scavenger (equal parts of 100 mM DTT, 1 M glucose, 20 mg/mL glucose oxidase, 4 mg/mL catalase, 100 mM MgCl_2_), 4.5 µl of KIFBP elution buffer (20 mM K-HEPES, 50 mM KCl, 1 mM DTT) or 4.5 µl of 1 µg/µl KIFBP, and lysate (volume determined experimentally). Block the chamber for 3 minutes with 15 mg/mL BSA/P12. Flow in the motility mix, seal the chamber with melted wax, and image immediately.

### Microscopy

Fixed and live cell images of N2A-A5 cells were acquired on a Ti-2E inverted microscope (Nikon Instruments) both driven by NIS Elements Software (Nikon Instruments). Images were captured using either a Clara cooled charge-coupled device (CCD) camera (Andor) or Prime BSI scientific complementary metal-oxide-semiconductor (sCMOS) camera (Teledyne Photometrics) with a Spectra-X light engine (Lumencore). Imaging was conducted with a Plan Apo 20X 0.75 numerical aperture (NA). For live cell-imaging, cells were imaged in CO_2_-independent media (Gibco; 18045088) supplemented with either 10% or 1% fetal bovine serum (FBS, Gibco; 16000044) within environmental chambers held at 37°C. Live cell images were processed and analyzed using ImageJ/Fiji (Schindelin et al., 2012; Schneider et al., 2012).

Fixed cell images for analysis were acquired using the Agilent BioTek Cytation 5 cell imaging multimode reader with wide field of view camera outfitted with a 20X phase contrast objective (Agilent; 1320517) and the following filter cubes: DAPI (Agilent; 1225100) and GFP (Agilent; 1225101). Multichannel fluorescent images were captured at 20X magnification in widefield mode in a 5 X 5 image montage in the center of the well of the fixed 24 well glass bottom plate (Cellvis; P24-1.5H-N). Focus was achieved using fixed height plus offset and adjusted in each well accordingly. Image tiles were stitched to generate a single large image for neurite outgrowth analysis. Stitched images were background subtracted before analysis for optimal visualization of neuronal outgrowths.

### Neurite Outgrowth Analysis and Quantification

Fixed cell images acquired using the Agilent BioTek Cytation 5 Cell Imaging Multimode Reader were analyzed after image processing using the Agilent BioTek Gen5 Neurite Outgrowth Module. Cell number is measured and quantified using DAPI signal while neurite lengths are measured using tubulin signal. Average neurite lengths are quantified by dividing the total neurite length by the total number of cells. N2A-A5 cells are considered differentiated if there is >/= 1 neurite outgrowth connected to the cell body that is > 10 µm. Percent of differentiated cells is calculated by total number of differentiated cells divided by total number of cells. Each data point represents a measurement from an imaged field of view.

### Statistical Analyses

Statistical analysis was performed using GraphPad Prism (GraphPad Software; Version 10). Specific statistical tests for reported data are indicated in the figure legends. All data represent a minimum of three independent experiments.

### DATA AVAILABILITY

RNA-Seq has been deposited in NCBI Gene Expression Omnibus (GEO) (pending approval).

## ACKNOWLEDGMENTS

This work was supported by NIH R35 GM144133 to JS, NIH R35 GM131744 to KJV, NIH R01GM141119 to MC, and a Vermont Space Grant Consortium Graduate Student Fellowship to SCP. We acknowledge the University of Vermont Integrative Genomics Resource CORE Facility (RRID:SCR_021775) for RNA sequencing library preparation, the University of Vermont Bioinformatics Shared Resource CORE Facility (RRID:SCR_027493) for Bulk RNA sequencing analysis, and The Vermont Biomedical Research Network Proteomics Facility (RRID: SCR_018667) supported through NIH grant P20GM103449 for mass spectrometry sample analysis.

## Notes

### Competing Interest Statement

The authors have declared no competing interest.

