## Supplemental Figures for "A systems-level proteomic analysis identifies kinesin targets of KIFBP during neuronal development"

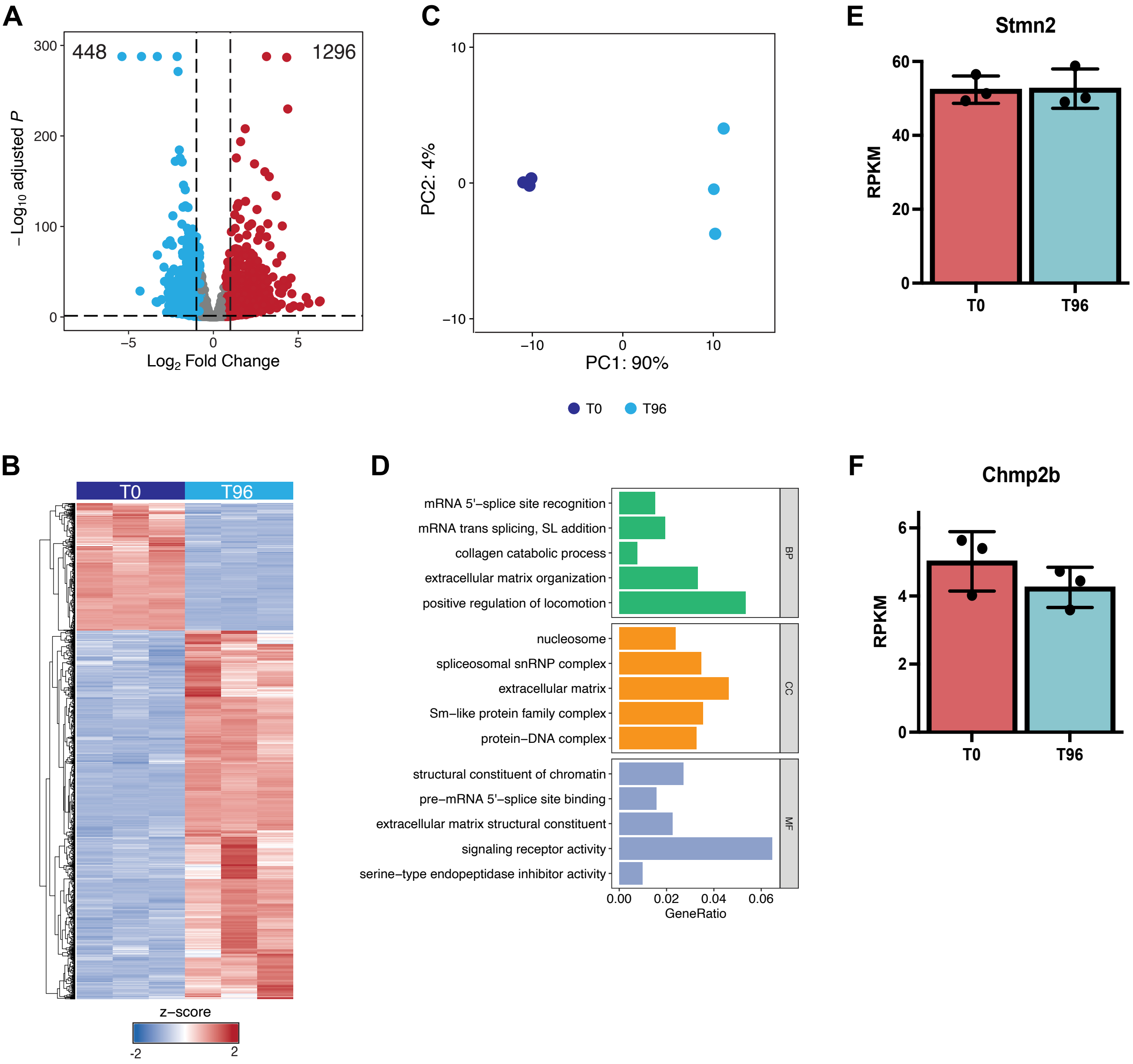


**Supplemental Figure 1**. Differentiation of N2A-A5 cells induces global gene expression changes. RNA sequencing was performed on dividing N2A-A5 cells at time 0 hour (T0) and differentiated cells after 96 hours (T96). (A) Volcano plots showing differentially expressed genes (DEG) between T96 vs T0 (FDR < 0.05, log2 fold change). (B) Hierarchical clustering of samples and DEG identified between T96 vs T0 (FDR < 0.05, log2 fold change). (C) Principal component analysis of the 500 most variable genes. Percentages in parentheses indicate the proportion of variance explained by each principal component. (D) Gene Ontologies enrichment analysis of DEGs including Biological Processes (BP), Cellular Components (CC), and Molecular Function (MF) categories. (E, F) Barplot of Stmn2 expression (E) and Chmp2b expression (F) between T0 and T96 shown in reads per kilobase per million (RPKM) normalized expression for cell types.

A

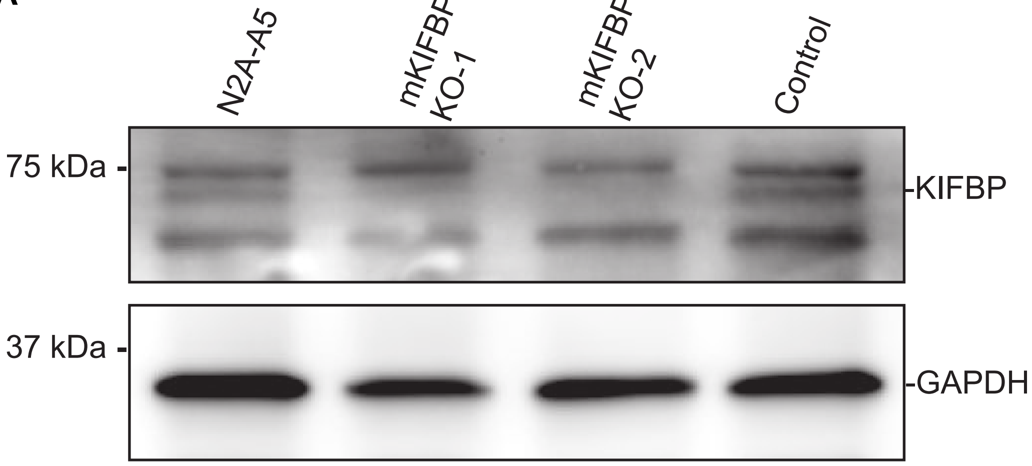
N2A-A5
mKIFBP KO-1
mKIFBP KO-2
Control
75 kDa
37 kDa
KIFBP
GAPDH
